# Modelling glioblastoma tumour-host cell interactions using adult brain organotypic slice co-culture

**DOI:** 10.1101/166967

**Authors:** Maria Angeles Marques-Torrejon, Ester Gangoso, Steven M. Pollard

## Abstract

Glioblastoma (GBM) is an aggressive incurable brain cancer. The cells that fuel the growth of tumours resemble neural stem cells found in the developing and adult mammalian forebrain. These are referred to as GBM stem cells (GSCs). Similar to neural stem cells, GSCs exhibit a variety of phenotypic states: dormant, quiescent, proliferative and differentiating. How environmental cues within the brain influence these distinct states is not well understood. Laboratory models of GBM tumours can be generated using either genetically engineered mouse models, or via intracranial transplantation of cultured tumour initiating cells (mouse or human). Unfortunately, these approaches are expensive, time-consuming, low-throughput and ill-suited for monitoring of live cell behaviours. Here we explored whole adult brain coronal organotypic slices as a complementary strategy to remove the experimental bottleneck. Mouse adult brain slices remain viable in a neural stem cell serum-free basal media for several weeks. GSCs can therefore be easily microinjected into specific anatomical sites *ex vivo*. We demonstrated distinct responses of engrafted GSCs to different microenvironments in the brain. Within the subependymal zone – one of the adult neural stem cell niches – a subset of injected tumour cells could effectively engraft and respond to endothelial niche signals. GSCs transplanted slices were treated with the anti-mitotic drug temozolomide as proof-of-principle of the utility in modelling responses to existing treatments. Thus, engraftment of mouse or human GSCs onto whole brain coronal organotypic brain slices provides a convenient experimental model for studies of GSC-host interactions and preclinical testing of candidate therapeutic agents.

## Introduction

Glioblastoma multiforme (GBM) is a highly aggressive malignant brain tumour. It is the most malignant form of glioma. Standard treatments involve combined surgery, radiotherapy, and adjuvant temozolomide (TMZ) chemotherapy (Stupp et al., 2005). However, long-term survival rates are extremely poor. Various obstacles hamper development of effective therapies, including: pervasive tumour cell infiltration, genetic heterogeneity (both intra- and inter-tumoural), therapeutic resistance, blood-brain barrier, and lack of biological understanding of the disease. Improved experimental models will help address some of these issues.

GBM tumour cells disseminate widely across many brain regions, often following neuronal tracts and vasculature. Cells are therefore exposed to diverse microenvironments, such as specific repertoires of cell matrix and growth factors, or cellular niches (e.g. perivascular, invasive or hypoxic). These environmental cues steer glioma stem cell (GSC) fate, affecting quiescence, proliferation, survival and differentiation pathways (Codrici et al., 2016; Gilbertson and Rich, 2007). Modelling these various tumour cell-host brain interactions is therefore vital for improved understanding of disease biology and development of new therapeutic strategies.

GSCs highjack many molecular programs that regulate neural stem cell self-renewal. Improved understanding of mechanisms controlling neural stem cell fate will therefore likely lead to new insights into the disease and identification of critical therapeutic targets. Neural stem cells (NSCs) are located within the lateral walls of the forebrain ventricles in a region known as the subependymal zone (SEZ) (Doetsch et al., 1999). The SEZ provides a specific niche that sustains the NSCs throughout life. NSCs are exposed to a myriad of cell-cell signals and ECM interactions that steer NSC fate, such as: endothelial cells, ependymal cells, and cerebral spinal fluid (Mirzadeh et al., 2008; Shen et al., 2008; Silva-Vargas et al., 2016). Understanding how tumour cells respond to normal SEZ is important, as this might serve as a reservoir of tumour cells that underlie relapse in some patients (Piccirillo et al., 2015).

Patient-derived GSCs can be routinely expanded *in vitro* using culture media developed for NSCs, either in suspension or adherent culture (Galli et al., 2004; Hemmati et al., 2003; Lee et al., 2006; Pollard et al., 2009; Xie et al.). Orthotopic transplantation of freshly isolated or cultured GSCs into the adult rodent brain using stereotaxic surgery is the ‘gold standard’ method to test tumour-initiating potential. However, animal surgery and transplantation deep into the brain provides limited experimental outputs: survival curves and end-point analysis of the resultant tumours. Typically, these experiments take weeks or months and are non-trivial to setup. They do not enable direct inspection of single cell behaviours such as invasion, monitoring of quiescence and differentiation or responses to genetic or chemical perturbations. These practical constraints have limited the scale and scope of studies aimed at understanding and treating gliomas. To address this we explored the utility of organotypic slice cultures to monitor GSC-host interactions.

Organotypic brain slice cultures were first developed in the 1960’s (Crain, 1966). Since then they have been widely used by neuroscientists, particularly in studies of neuronal function and circuits (reviewed in (Humpel, 2015)). Microdissected regions are cultured above a semipermeable membrane in a cell culture insert and exposed to serum-containing media from below. An example of their value are studies using hippocampal slices cultures, widely deployed for studies of synaptic plasticity and memory (Gahwiler et al., 1997). Organotypic slice cultures overcome some of the difficulties of *in vivo* studies as they provide *ex vivo* access to brain tissue architecture, while still enabling direct observation and cell manipulations in the culture dish (Humpel, 2015).

In this study we demonstrate improved conditions enabling serum-free culture of adult coronal whole brain slices in a manner that enables tracking of GSC behaviors over several weeks. Our experimental approach provides a useful new strategy to explore GBM. This model bridges the ‘experimental gap’ between *in vitro* cell culture models and *in vivo* orthotopic transplantations. As an exemplar of the utility of this approach we confirm engraftment of GSCs around blood vessels in the slice culture and demonstrate how the method can be used in preclinical studies of anticancer agents.

## Results

### Whole adult brain coronal slice cultures are viable for weeks in serum-free neural stem cell media

Most studies employing organotypic slice cultures use postnatal mice and dissect specific brain regions (e.g. hippocampus). However, GBM is predominantly a disease of adults and cells disseminate across all brain regions. We therefore focused on whole brain slices, reasoning that even short-term viability, for days or weeks, could provide a useful model for testing tumour cell-host brain interactions.

Adult brains were harvested from young adult mice (~4 weeks) and the olfactory bulbs and cerebellum were removed (Figure 1A and B). We generated whole-brain coronal sections using a vibratome to cut ~200um slices at the level of the forebrain ventricle (6 slices per brain). Each section was placed on to a semi-permeable membrane culture insert and cultured in a six well cell culture plate (Figure 1B).

**Figure 1.**
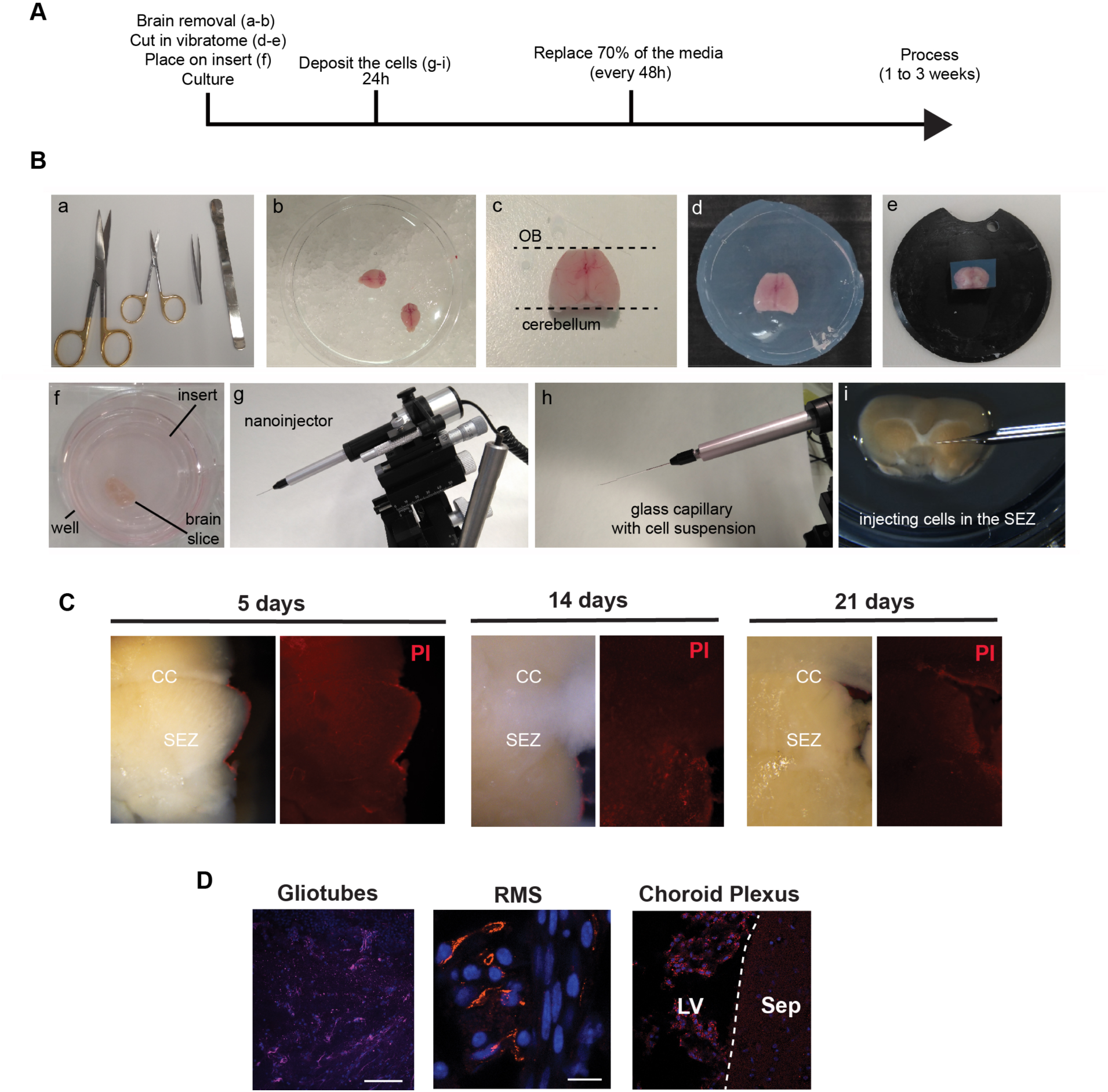
Overall experimental strategy and tissue processing. (A) Summary of experimental procedure to generate slices. (B) Experimental steps in the harvesting, mounting, slicing and injection of brain tissue; (a) scissors, forceps and spatula are used to isolate and dissect the whole brain; (b) Image of whole adult mice brain on ice following harvesting; (c) Dorsal image of whole brain following removal of the olfactory bulb (OB) and cerebellum; (d) Embedded brain in low melting agarose; (e) Attached brain to the support of the vibratome; (f) ~250μm coronal brain slice of placed onto a cell culture insert in a 6 well plate with neural stem cell basal media; (g) nanoinjector mounted on a micromanipulator used for injection of small volumes of cells, with mounted glass capillary (h), containing cell suspension; and (i) microinjection of cells into the SEZ of a coronal brain slice on the cell culture insert. (C) Viability of the tissue assessed using propidium iodide after 5, 14 and 21 days. (D) Immunocytochemistry following 7 days slice co-culture for: GFAP positive gliotubes (magenta; left panel); DCX-positive neuroblasts (middle panel) and choroid plexus (autofluorescence; right). Nuclear counterstaining with DAPI in each (blue). SEZ: subependymal zone; CC, corpus callosum; LV; lateral ventricle, Sep: Septum. RMS; Rostral migratory stream. Scale bars: 100μm (left) and 10μm (right).

Established organotypic brain slice protocols require high levels of serum or growth factors. However, serum exposure will trigger astrocyte differentiation of NS cells (Conti et al., 2005) (Figure S1A-B). Culture media lacking serum or exogenous growth factors was therefore tested. This reduces the risk of cell fate being primarily directed by additives in the culture media, rather than endogenous tissue-derived signals. Dying cells were identified at edges of the dissected region, using staining for propidium iodide (PI). Serum-free NS cell-permissive culture media could indeed support slices over several weeks (Figure 1C). By contrast, in basal media with no N2 or B27 supplements, tissues became necrotic in days (Figure S1C).

Within the healthy coronal sections, we were able to detect Dcx-expressing neuroblasts, gliotubes and choroid plexus, suggesting the tissue retained key features of a viable neurogenic niche (Figure 1D). In summary, whole brain coronal slices are viable for several weeks in serum-free media – much longer than we anticipated – thereby providing an opportunity to assess responses of transplanted GSCs.

### Patient-derived glioblastoma stem cells engraft into the mouse SEZ and retain expression of quiescent NSC markers CD133 and CD9

We next tested the potential of patient-derived human GSCs to engraft into the slices. G7-GFP cells have previously been shown to be highly invasive when transplanted into striatum of immunocompromised mice (Stricker et al., 2013). We first tested microinjection of 10000 cells into the SEZ (Figure 2A). One week later using live cell imaging we noted large numbers of healthy GFP expressing cells successfully engrafted. After 2 weeks slices were fixed and immunocytochemistry (ICC) confirmed cells were viable and based on Ki67 and Stem121 expression ~10% were actively proliferating (Figure 2B). We next assessed the known cancer stem cell marker CD133 in the GFP cells after three weeks. Interestingly, CD133 expressing cells were identified in a subset of cells, suggesting that GSCs generate phenotypic heterogeneity in the slices and retain cells with cancer stem cell phenotype (Figure 2C).

**Figure 2.**
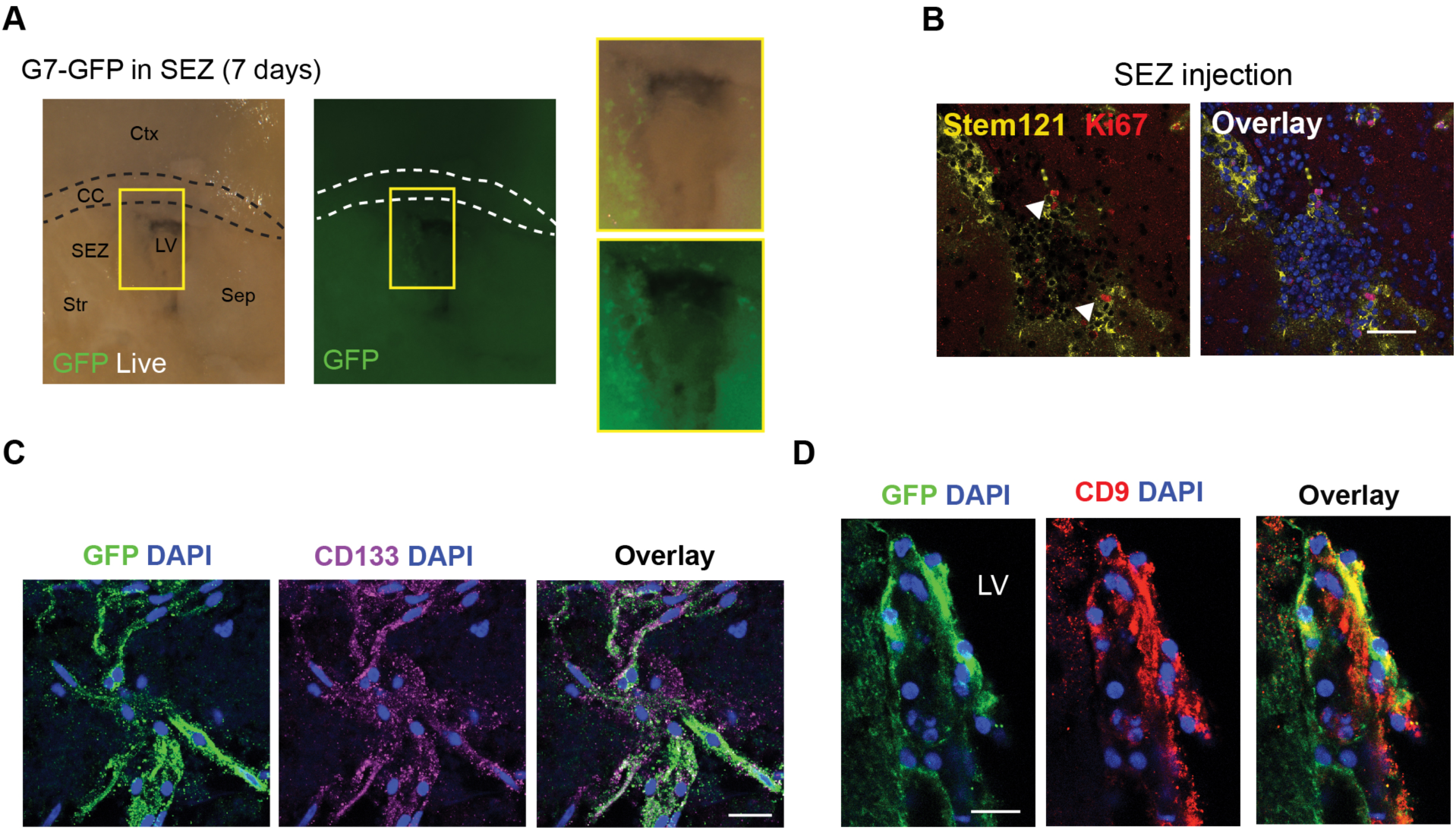
Human patient-derived glioma stem cells engraft in adult mouse subependymal zone. (A) Direct microinjection of human line into adult SEZ and visualization of engrafted live cells using a constitutive GFP reporter. Right panels are zoom of left panels. (B) Immunostaining for human specific cytoplasmic antigen (Stem121; yellow) and Ki67 (red) after 14 days. (C and D) Marker analysis after 21 days of ex vivo culture; (C) immunocytochemistry for CD133 (purple); (D) CD9 (red). Nuclear counterstaining with DAPI (blue). Scale bars: B = 50 μm and C, D = 20 μm.

New molecular markers associated with quiescent NSCs have recently been uncovered using single cell RNA-seq approaches (Llorens-Bobadilla et al., 2015; Shin et al., 2015). The transmembrane glycoprotein tetraspin (CD9) was identified as a putative marker of qNSCs. Many of the G7 cells that were transplanted within the SEZ expressed CD9 (Figure 2D). We conclude that human GSCs can engraft, survive and proliferate in whole-brain slice co-cultures for at least 3 weeks, and can retain expression of key cancer stem cell markers.

### Patient-derived glioblastoma stem cells have distinct responses to region-specific adult brain microenvironments

We next compared how cells would respond in the SEZ versus other brain regions in terms of their proliferation and differentiation. Four distinct regions were tested: striatum, corpus callosum (CC), cortex and SEZ (Figure 3A). One week after microinjection into a lateral region of the CC, we noted many G7 cells aligning with and dispersing across the while matter tracts and displaying infiltrative morphology. These cells displayed reduced Ki67 and increased GFAP compared to cells deposited in parallel within the SEZ of the same slice (Figure 3B-C). Similar findings were seen for cells within the cortex and striatum. Thus, slice cultures provide a convenient method to quickly assess responses of GSCs to the diverse anatomical microenvironments within the adult brain. This enables future exploration of various candidate factors that might influence cell fate within the SEZ niche.

**Figure 3.**
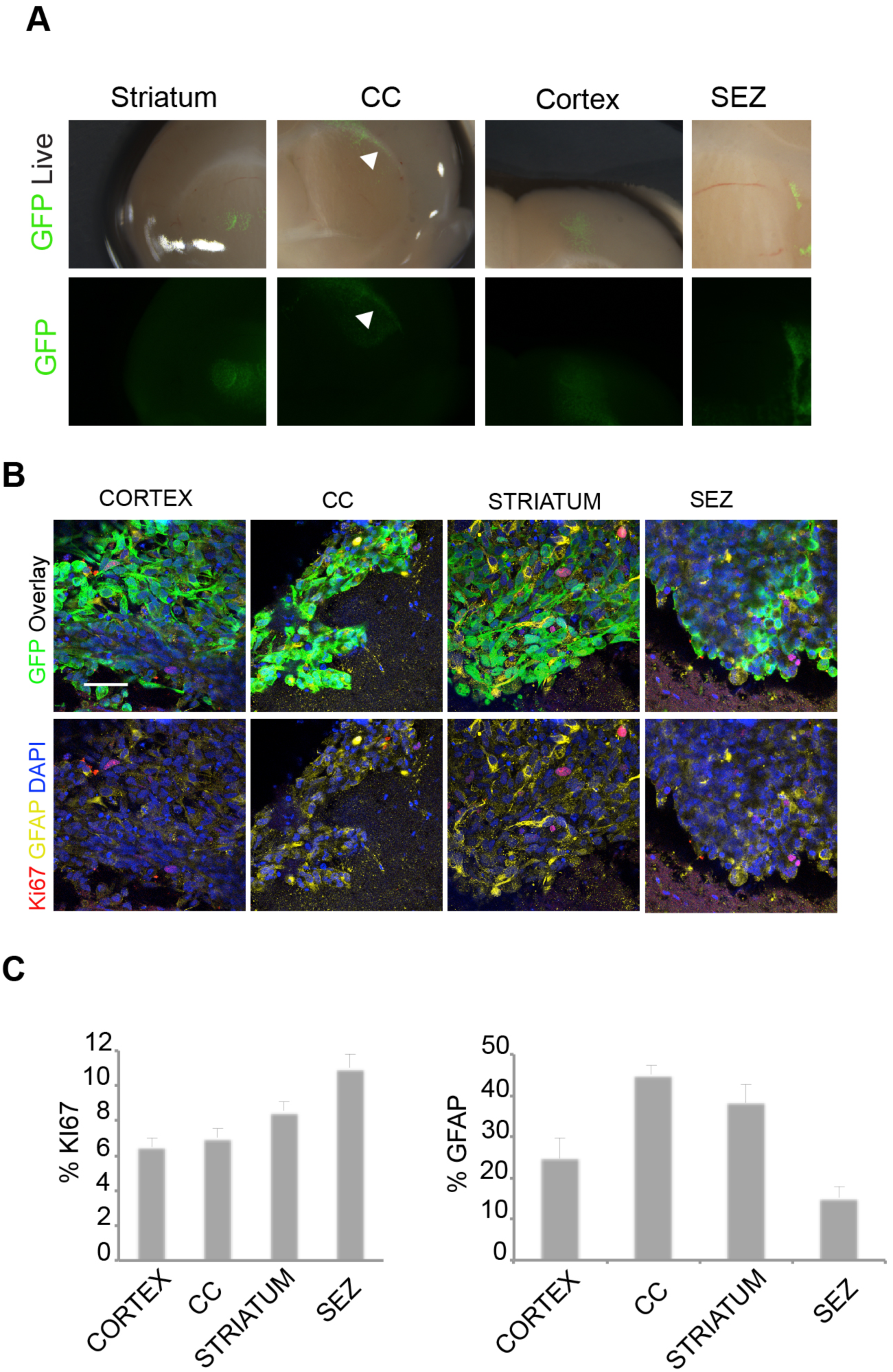
Differential proliferative responses to human glioma stem cells engrafted into different brain regions. (A) Live images of G7-GFP human glioma stem cells deposited into distinct regions of the same coronal brain slice after 7 days. (B) Immunocytochemistry for GFAP (yellow) and Ki67 (red) and GFP (green). CC: corpus callosum. (C) Quantification of the percentage of proliferating and differentiating cells (Ki67 and GFAP, respectively).

### Mouse glioblastoma-initiating cells engraft into the subependymal zone and can juxtapose to endothelial cells

To minimize disruption of the niche and to enable injection of smaller volumes/numbers of cells (~100 cells in ~40nl) we performed transplantation of cells using a glass capillary linked to a microinjection pump (Figure 1B-C). This enabled more precise and localized injection into the walls of the lateral ventricle (Figure 4A). We used a previously characterised mouse tumour-initiating cell line, termed IENSGFP (Ink4A/Arf^−/−^ deleted with EGFRvIII overexpression) (Bruggeman et al., 2007). These cells stably express GFP from a constitutive promoter. In vitro they express GSC markers such as Nestin, Sox2, Olig2 (Figure S2A). IENS cells generate aggressive infiltrative tumours when transplanted *in vivo* (n=4) (Figure S2B). These were preferred to human patient-derived G7-GFP, as these displayed brighter GFP and were less clumpy.

**Figure 4.**
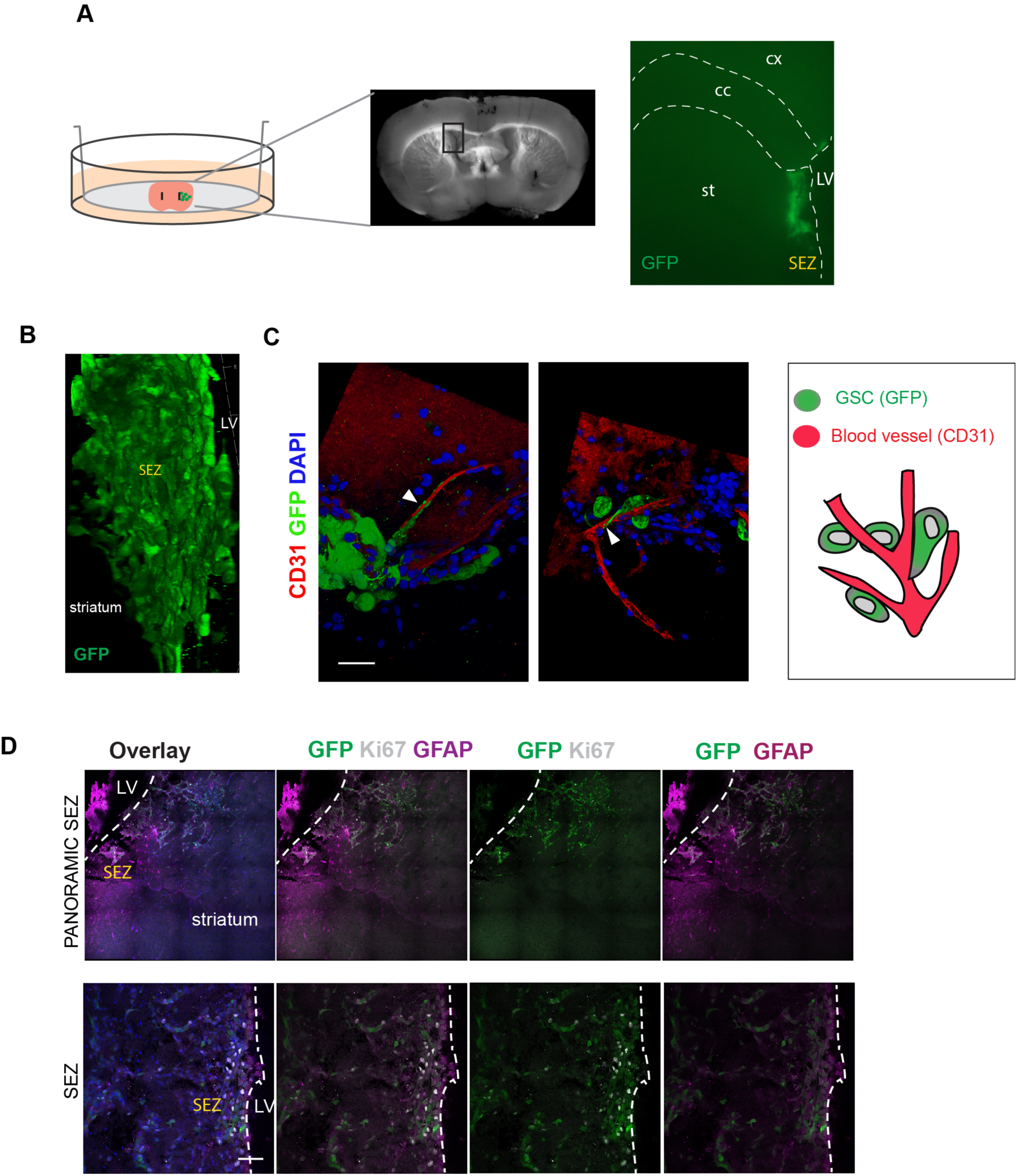
Glioma stem cell (GSC) in the SEZ niche. (A) Left, Schematic of the experimental setup: co-culture of the GSC and the whole brain coronal sections. Right, panoramic position of the GSC (green) in the SEZ after deposition. (B) 3D image showing the engraftment and infiltration of the IENS-GFP in the brain slice after 5 days. (C) 3D image with immunostaining of the IENS-GFP contacting with the blood vessels. GFP (green), CD31 (red) and nuclear counterstaining with DAPI (blue). Schematic of the interaction GSC with brain blood vessels (right). (D) Top panels: Panoramic view of IENS-GFP cells in SEZ. Immunostaining for GFP (green), Ki67 (white), GFAP (purple) and nuclear counterstaining with DAPI (blue). Lower panels: Cells proliferating and expressing some GSC and astrocytic marker as GFAP and the proliferative marker Ki67. In (A) cx, cortex; cc, corpus callosum; st, striatum; SEZ, subependymal zone; LV, lateral ventricle. Scale in (C) 20μm (D) 50μm.

IENS cells were injected precisely into the SEZ through the wall of the lateral ventricle. Initially, they displayed a remarkably specific localization and even distribution throughout the SEZ (Supplementary Movie 1). Cells injected in parallel into the striatum also survived and engrafted, but dispersed locally (Figure S2C). This may suggest a preference, or homing, to the SEZ (Figure 4A), similar to previous reports for normal NSCs (Kokovay et al., 2010). When imaged using confocal microscopy we noted juxtaposition of GSCs with endothelial cells (Figure 4C and Supplementary movie 2). 5 days after microinjection the rate of proliferation was around 40-50% (measured as Ki67^+^ cells of the GFP cells). ~15% began to express high levels of the astrocytic marker GFAP (Figure 4D). Cells remained viable for two weeks and showed evidence of proliferation and local infiltration into surrounding regions (n=3) (Figure 4B and Movie 1 and 2).

### Glioblastoma stem cells engrafted into brain slices respond to the cytostatic effects of temozolomide

Whole brain slice cultures with engrafted GSCs provides a convenient, easier and higher throughput experimental system to explore the effects of pharmacological agents in than use of live whole animal studies. To demonstrate proof-of-principle of potential use in preclinical studies we explored effects of anti-mitotic treatments. Ara-C or Temozolomide (TMZ) have been widely used to explore neural stem cell behavior during regeneration and repair (Daynac et al., 2016; Doetsch et al., 1999). TMZ is the standard chemotherapy given to many patients with GBM. Both agents drive DNA damage and disrupt proliferation of tumour cells. TMZ is a DNA alkylating agent that often induces G2/M arrest.

Cellular responses were scored using immunocytochemistry for two markers: gamma-H2AX for double strand breaks, and p53 as an indicator of DNA damage response (Figure 5A). We first tested activity of each factor at previously reported effective doses on IENS-GFP cultures *in vitro*, TMZ at 1,10 and 100μM and AraC at 1 and 2 μM (Figure 5A-B). We next treated slices harbouring successfully engrafted IENS-GFP cells after 3 days with 100μM TMZ or 1μM Ara-C (24hr). Slices were then assessed for Ki67 and pHH3 using immunocytochemistry (Figure 5E-F). In each condition we observed a reduction in Ki67 and GFP double positive cells (100μM TMZ: 8%, 1μM Ara-C: 11%). Untreated control slices retained ~30% double positive (Figure 5E-F). Cytostatic responses of tumour cells to drug treatment can therefore easily be monitored within slice co-cultures.

**Figure 5.**
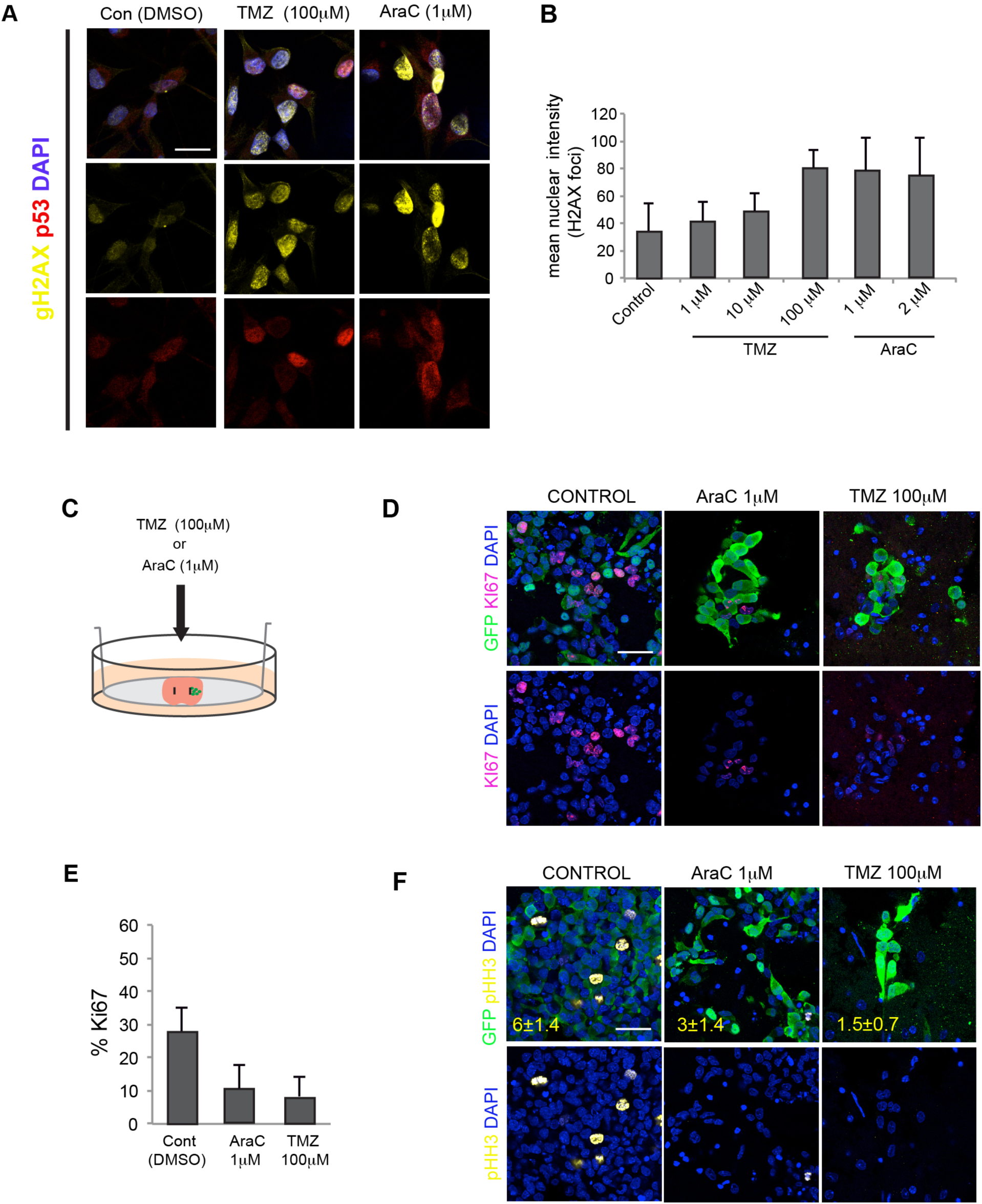
Temozolomide and Ara-C treatment of slice co-cultures. (A) DNA damage responses of mouse IENS-GFP cells following treatment various does of antimitotic agents AraC and TMZ. Immunocytochemistry for γH2AX (yellow) and p53 (red). Nuclear counterstaining DAPI (blue). (B) Quantitation of mean intensity of the nuclear H2AX foci (arbitrary units). (C) Schematic of the co-culture of IENS-GFP cells in the brain slices adding pharmacological inhibitors of the proliferation. (D) Immunostaining for GFP (green) and Ki67 (magenta) with DAPI (blue) for nuclear counterstaining. (E) Quantification of the percentage of Ki67 (proliferative cells) using the proliferative inhibitors in IENS-GFP after 24 hours. (F) Immunocytochemistry for GFP (green) and pHH3 (mitotic cells) (yellow) in IENS-GFP after treatments. Percentage of pHH3 positive cells are in red typing. Scale bar in (A) 10 μm (D), (F) 20μm.

## Discussion

Primary cell cultures of human GBM stem cells are an important and disease-relevant in vitro experimental tool. However, an obvious limitation of dissociated cell cultures is the difficulty in modelling interactions with the complex tissue environment. Here we have demonstrated that brain tumour cell interactions with host brain tissue can be explored effectively using co-culture of in vitro expanded mouse and human GSCs with adult brain slices.

Past organotypic methods have typically used micro-dissected regions of postnatal brain slices cultured in the presence of serum, as this facilitates the support of long-term viability (months) (Ullrich et al., 2011). Yet our observations indicate that whole brain adult coronal slices are viable in serum free media for several weeks. This is long enough to permit tracking of tumour-host tissue interactions, such as interactions with endogenous stem cell niches or white matter tracts and enabled us to test GSCs interactions with brain tissue. These methods therefore provide a tractable ex vivo model system that can now be exploited in both basic and translational studies of GBM. It is an approach that reduces the need for laborious and expensive mouse breeding or stereotaxic surgery, thereby increasing the speed and experimental throughput.

Avoiding exposure of transplanted cells to high levels of serum within the slice cultures enables a more physiological signaling environment to be maintained. This reduces the degree of serum-induced astrocyte differentiation which has hampered our previous studies (unpublished observations). Although serum free media has been used to maintain whole-mount tissue explants of the mouse SEZ for up to 16 hours (Kokovay et al., 2010), to our knowledge longer term survival of whole coronal brain slices in serum free media has not be reported for studies of GSCs. Surprisingly, we found that serum wasn’t needed in order to maintain healthy slices of the whole adult coronal brain for many weeks. Slices seems viable in the basal neural media with no exogenous growth factors and supplemented only with N2 and B27 hormonal supplements.

With viable coronal adult brain slices we were able to assess the responses of both mouse and human GSCs to distinct anatomical regions over several weeks of co-culture. As each mouse can provide up to 5 or 6 slices and cells can easily be injected in a spatially-restricted manner. Live cell imaging can be performed to track dynamic cell behaviors, such as interactions with blood vessels, infiltration or division (Supplementary Movie 1). For example, GBM cells infiltrate widely, and have been shown to use both neuronal tracts and blood vessels as a substrate and guide for migration (Farin et al., 2006; Krusche et al., 2016). The slice cultures reported here should be useful in probing such distinct cellular mechanisms of infiltration. Another future potential application will be the tracking of cell lineage reporters, especially with the advent of genome editing tools that can be applied in GBM (Bressan et al., 2017). This will also help shed light on mechanisms of GSC dormancy and quiescence. Future studies might also make use of short-term immunological responses to the engrafted tumour cells, such as interactions/activation of microglia.

We were particularly interested in assessing tumour cell responses within the SEZ. The SEZ provides a specific niche that sustains the NSCs, and a repertoire of cell-cell signals and ECM interactions that steer NSC fate, including: endothelial cells, ependymal cells, and cerebral spinal fluid (produced by the choroid plexus) (Mirzadeh et al., 2008; Shen et al., 2008; Silva-Vargas et al., 2016). We noted key elements of the healthy SEZ microenvironment were viable – ventricle, gliotubes, RMS, ependymal cells. Importantly, endothelial cells within this region are thought to serve as an important niche signal; indeed, we noted close interactions between vessels and tumour cells, with extended processes and wrapping around the vessel, highly reminiscent of normal NSC interactions (Kokovay et al., 2010).

We demonstrated that human GSCs can engraft effectively into the tissue of SEZ of immunocompetent. CD133 is expressed by many glioma stem cells. We observed expression of this marker in both the mouse and human GSCs. Another more recently proposed marker of the quiescent astrocytes is CD9. We and others have recently found increased levels of CD9 in GSCs compared to normal NSCs (Okawa et al., 2016b; Podergajs et al., 2015). CD9 is expressed on cells within the SEZ, suggesting ‘stemness’ may be effectively retained. As anticipated, reduced proliferative responses were noted when cells were deposited at other anatomical sites, such as corpus callosum. Distinct brain regions clearly will have significant differences in their ability to influence tumour cell proliferation and differentiation and this can now be further explored using this model.

A limitation of this technique might be the difficulties of achieving viable slice cultures past 3 weeks. Although we did note some loss of some tissue integrity past three weeks, we did not specifically assess later time points or search for modified culture regimes. This might be important to resolve in future, particularly for slower growing human GSCs, which take weeks to months to start initiate tumour growth in live xenografted mice.

A multitude of new therapeutic agents are emerging that will require effective preclinical studies before entering clinical trials for GBM. There is bottleneck and huge cost associated with testing of new pharmacological agents in living animals. The methods outlined here offer one potential solution. To demonstrate potential utility as preclinical model, we tested the responses of cells to anti-mitotics (AraC and TMZ). TMZ is used in many GBM patients, yet our understanding of how it influences distinct compartments of the GSC and resistance mechanisms remains limited. Future candidate drugs will need to be explored alongside TMZ to search for the most effective doses and regimes.

In conclusion, the organotypic method presented here provides a simplified model for assessing responses of GSCs to various brain anatomical sites and microenvironments. This experimental model will therefore complement existing in vitro and in vivo models, helping to prioritise genes and pathways controlling key malignant properties of GBM and aiding the preclinical testing of new anti-cancer agents.

## Materials and Methods

### Organotypic adult brain slice culture

4-8 week-old C57BL/6 mice were used. More consistent results were often obtained using the younger animals – particularly the viability after 2-3 weeks culture. The brain was removed from the skull and transferred to a 10cm^2^ tissue culture dish with sterile PBS and placed on ice (Figure 1A-1B). Cerebellum and olfactory bulb were removed (Figure 1B) and remaining forebrain transferred into a 35mm^2^ dish with pre-warmed 3% SeaPlaque™ agarose (50100 Lonza) (Figure 1A-1B). Upon cooling in ice the block was removed and cut using a scapel into a ~ 2cm cube around the brain. Prior to sectioning a 6-well plate was prepared: in each well we introduced one cell culture insert (PICMORG50 Millicell) and added bellow it 1mL of culture media in serum-free basal NS cell media, DMEM:F12 supplemented with N2 and B27 (Life Technologies). The embedded brain was placed in the circular vibratome plate with adhesive (Figure 1A-1B). The vibratome (Leica VT1000 S) plate was fixed in the platform and was filled with PBS with penicillin-streptomycin (Gibco 15140-122 1:100). 250 μm thick slices were cut, with vibrating frequency set to 8 and speed to 3. Each slice was transferred using a small brush onto the top of a Millipore culture insert (Figure 1A-1B). Six slices were cut per animal along the SEZ. The platform has to be maintained cooled all the time. We obtained six slices around the forebrain ventricle. The 6 well plate was placed in an incubator at 37ºC + 5% CO_2._ Slices were incubated for 1-2 days prior to cell transplantation.

### Glioblastoma cancer stem cell transplantation onto brain slices

G7 human GBM stem cell cultures have been previously described (Stricker et al., 2013). For human cell transplants a standard Gilson pipette was used to deposit 0.2 ul of cells onto the centre of the striatum, the typical injection site when performing stereotaxic surgery for tumour initiation assays. These cells engrafted well into the slice and their integration could be observed the following day.

IENS-GFP mouse cells were a gift from Prof M. Lohuizen (Bruggeman et al., 2007). Both mouse and human GNS cells were grown using conditions previously described (Pollard et al., 2009). After centrifugation cells were harvested and resuspended in media at 100,000 cells/ul. Cells were used immediately for transplantation in the SEZ. Two different methods of cells injection were used. A manual using a p2 Gilson pipette was used to injected 0.5ul, while an auto-nanoliter injector (Nanoject II, Drummond Scientific Company) was used for 40-100nl injections. For the injector we use glass capillaries pulled using an automated needle puller (tip diameter, 10–20 μm; Drummond) (Figure 1B). 4000 cells were transplanted per injection. 20000 cells were transplanted when the P2 pipette was used. To facilitate engraftment and prevent wide dispersion of the 0.2 ul of cells, we used forceps to make a small indentation in the transplant site prior to delivery of the cells. Slices with cells were incubated at 37ºC + 5% CO_2_ for 7 days, with fresh NS cell media (no EGF or FGF-2) added every 2 days. Cell engraftment was observed 4d after transplantation and cells could were monitored using a fluorescence stereomicroscope (Leica M165 FC). For anti-mitotic treatments in the IENS-GFP in vitro, different doses of the cytosine arabinoside (AraC) (Sigma) were added to the complete media. For anti-mitotic treatments of IENS-GFP in the slices, the cells were grown in the brain slice for 3 days and the AraC was added to the complete media at different doses at 1uM and TMZ at 100uM for 24hr.

### Immunocytochemistry

Media was removed and exchanged for 1mL of freshly prepare 4% paraformaldehyde (PFA). 1-2 mL was also placed gently on top to cover the slice. After 2 hours PFA was removed and brain slices were wash with PBS three times. Slices were transfer using a brush to a 24 well plate. Slices were incubated at room temperature 1.5hr in blocking solution (0.2% Triton X-100 and 3% Goat Serum; Sigma). Primary antibodies GFAP (G3893 Sigma 1:100), Ki-67 (RM-9106 Thermo Fisher 1:100), GFP (13970 Abcam 1:100), CD31 (14-0311-81 eBiosience 1:100), CD9 (14-0091-82, eBioscience), CD133 (MAB4310, Millipore), DCX (AB2253 Millipore), Nestin (1/10 Hybridoma Rat 401, DSHB), stem121(440410, Cellalartis), pHH3 (50-9124-41, eBioscience), H2AX (phosphor S139) (ab81299, Abcam).

For H2AX staining cells were fixed in methanol-acetone 1:1 for 10min at room temperature. Positive cells were scored using the Fiji image analysis software. For IHC, the primary antibodies were incubated for 2d at 4ºC. After three washes with PBS, slices were incubated with appropriate Alexa Fluor (Life technologies) secondary antibody and DAPI (D9542-SIGMA) with for 4-6 hours. Slices were washed three times and were mounting in a slide and immersed in FluoroSave^TM^ Reagent (345789 Calbiochem). Slices were examined with a confocal microscope (Leica TCS SP8). Propidium iodide (14289-25 CAYMAN) was used at 5ug/ml in PBS for 5 min and the tissue was analyzed under the fluorescent steromicroscope.

## Acknowledgements

We are very grateful for the support provided for imaging from Bertrand Vernay (University of Edinburgh). Raul Bressan and Kirsty Ferguson provided helpful comments on the manuscript. MAMT was supported by a Postdoc EMBO Long-Term Fellowship. EG was supported by a Postdoc fellowship from Fundacion Ramon Areces (Spain). SMP is supported by a Cancer Research UK Senior Research Fellowship (A19778).

---

**Movie 1** (for Figure 2). Living IENS-GFP moving thought the brain tissue after 10 days.

**Movie 2** (for Figure 2). 3D reconstitution of the IENS-GFP (green) engraftment in the SEZ after 5 days.

**Movie 3** (for Figure 2). 3D reconstitution of reconstitution showing the interaction between the IENS-GFP (green) and blood brain vessels (red). Nuclear counterstaining with DAPI (blue or grey).

**Supplementary Figure 1.**
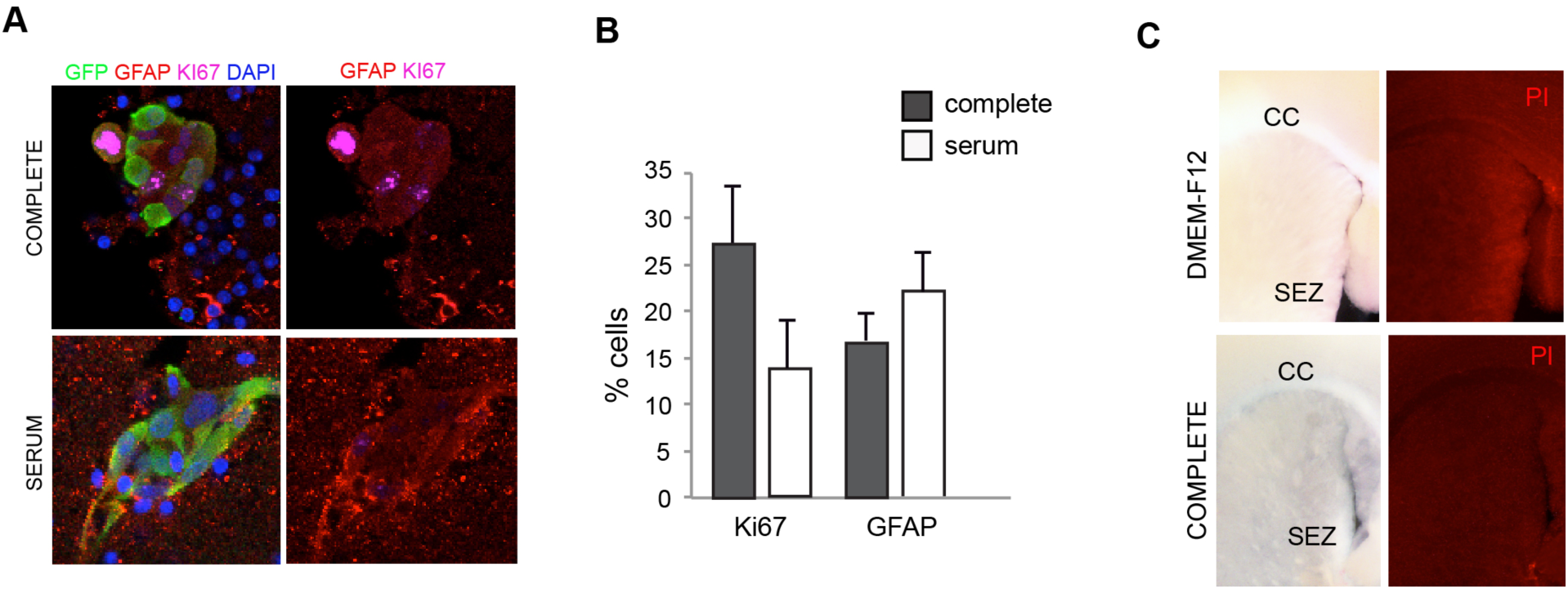
(A) Immunostaining for GFAP (red) and Ki67 (magenta) of IENS-GFP cells deposited in the brain slices with complete media with/without serum. (B) Quantification of the percentage of cells positive for GFAP and Ki67. (C) PI staining in brain slices after 24 hours. Slices were cultured with DMEM-F12 or with complete media.

**Supplementary Figure 2.**
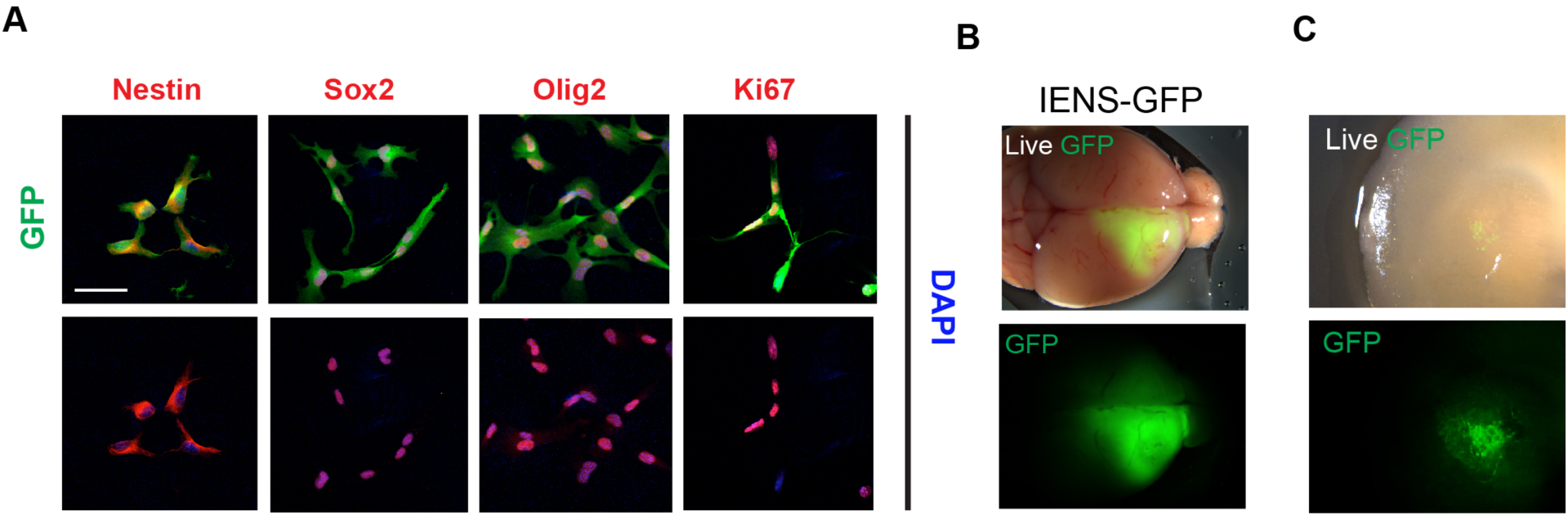
(A) IENS-GFP (green) express GSC markers Nestin, Sox2, Olig2 and Ki67 (red) with nuclear counterstaining DAPI (blue) and (B) are tumorigenic. (C) IENS-GFP deposited in the striatum. Scale bar in A: 20 μm.

